# Endothelial TGF-β signaling instructs smooth muscle development in the cardiac outflow tract

**DOI:** 10.1101/2020.01.30.925412

**Authors:** Giulia L.M. Boezio, Anabela Bensimon-Brito, Janett Piesker, Stefan Guenther, Christian S.M. Helker, Didier Y.R. Stainier

## Abstract

The development of the cardiac outflow tract (OFT), which connects the heart to the great arteries, relies on a complex crosstalk between endothelial (ECs) and smooth muscle (SMCs) cells. Defects in OFT development can lead to severe malformations, including aortic aneurysms, which have often been associated with impaired TGF-β signaling. To further investigate the role of TGF-β signaling in OFT formation, we generated zebrafish lacking the type I TGF-β receptor Alk5 and found a strikingly specific dilation of the OFT. *alk5* mutants also exhibit increased EC numbers, extracellular matrix (ECM) and SMC disorganization. Surprisingly, endothelial-specific *alk5* overexpression in *alk5* mutants rescues both endothelial and SMC defects. Furthermore, modulation of the ECM gene *fibulin-5*, a TGF-β target, partially restores OFT morphology and function. These findings reveal a new requirement for endothelial TGF-β signaling in OFT morphogenesis and suggest an important role for the endothelium in the etiology of aortic malformations.

## Introduction

The cardiovascular system is essential to deliver blood to the entire organism. Within the heart, however, the high pressure deriving from ventricular contractions needs to be buffered to avoid damage to the connecting vessels. The cardiac outflow tract (OFT), located at the arterial pole of the heart, fulfills this role and is a vital conduit between the heart and the vascular network (Kelly and Buckingham, 2002; Sugishita et al., 2004). Its importance is confirmed by the fact that errors in OFT morphogenesis lead to almost 30% of all congenital heart defects (CHD) in humans (Neeb et al., 2013). These malformations include defects in the alignment and septation of the OFT, such as persistent truncus arteriosus (PTA), transposition of the great arteries (TGA) and overriding aorta (OA), as well as coarctation or dilation of the arteries exiting the heart (Neeb et al., 2013; Anderson et al., 2016). However, the etiology of these malformations remains unclear, due to their multifactorial causes and the complex interplay between the different cell types in the developing heart.

The development of the OFT starts with the formation of a simple tube lined by endothelial cells (ECs) and surrounded by myocardium, both derived from late-differentiating second heart field (SHF) progenitors (Kelly and Buckingham, 2002; Paffett-Lugassy et al., 2017; Felker et al., 2018). Later, this tube switches from a myocardial to an arterial phenotype and becomes surrounded by smooth muscle cells (SMCs) of SHF and neural crest origins. In mammals, the OFT undergoes septation, cushion formation, and rotation, giving rise to the final mature structure that forms the trunk of the aortic and pulmonary arteries. In other vertebrates like zebrafish, in which the cardiac chambers do not undergo septation, the OFT does not divide and forms what is considered a “third chamber” – the bulbus arteriosus (Grimes and Kirby, 2009; Zhou et al., 2011; Guner-Ataman et al., 2013; Knight and Yelon, 2016; Paffett-Lugassy et al., 2017). Considering the cellular contributions to OFT development, most of the attention has been focused on the SMCs, cardiac neural crest, and cardiomyocytes (CMs) (Kelly and Buckingham, 2002; Buckingham et al., 2005; Waldo et al., 2005a; Waldo et al., 2005b). However, the endothelial lining of the OFT is also likely to play major roles.

Multiple signaling pathways including BMP, Notch, FGF, Wnt, and TGF-β have been implicated in OFT development (Neeb et al., 2013). In particular, mutations in several of the *TGF-β* family members have been associated with severe congenital heart diseases, such as PTA and aneurysm of the great vessels (Todorovic et al., 2007; Gillis et al., 2013; Takeda et al., 2018). Despite the clear importance of the TGF-β signaling pathway in the development and homeostasis of the OFT and connecting vessels, the molecular mechanisms underlying these defects remain elusive, due to the context-dependent and controversial role of this pathway (Massague, 2012; Cunha et al., 2017; Goumans and Ten Dijke, 2018; Zhang, 2018).

Activin receptor-like kinase 5 (Alk5, aka Tgfbr1) is the main type I receptor of the TGF-β signaling pathway. In mouse, *Alk5* expression is present specifically in the great arteries and the heart, mostly enriched in the SMC layer of the aorta (Seki et al., 2006). A global mutation of *Alk5* in mouse leads to embryonic lethality due to brain hemorrhage and presumed cardiac insufficiency (Carvalho et al., 2007). Notably, these phenotypes are recapitulated by the EC-specific deletion of *Alk5* (Sridurongrit et al., 2008). Conversely, despite its reported expression pattern, loss of *Alk5* in mouse cardiomyocytes, pericytes or SMCs does not lead to any obvious defect during the development of the heart or great vessels (Sridurongrit et al., 2008; Dave et al., 2018).

Despite the evidence for a potential role for ALK5 in the endothelium, most studies in OFT and aortic pathologies, such as aortic aneurysms, have been focused on the role of TGF-β signaling in SMCs and not ECs (Choudhary et al., 2009; Guo and Chen, 2012; Gillis et al., 2013; Yang et al., 2016; Takeda et al., 2018). In particular, aneurysms have been described as a weakening of the aortic wall due to SMC-specific defects, resulting in the dissection of the vessel (Takeda et al., 2018). Only recently have a few studies started considering the endothelium as a potential therapeutic target in OFT and aortic pathologies (van de Pol et al., 2017; Sun et al., 2018). The close proximity of ECs with SMCs makes their cross-talk essential for aortic development and homeostasis (Lilly, 2014; Stratman et al., 2017; Segers et al., 2018). However, the early lethality of the *Alk5* global and EC-specific KO animals prevents a deeper investigation of the role of this gene in heart and OFT development.

The use of zebrafish as a model system can help overcome the issues associated with early embryonic lethality resulting from severe cardiovascular defects (Stainier and Fishman, 1994). Moreover, this system allows a detailed *in vivo* analysis of the phenotype and the assessment of cardiovascular function during embryogenesis thanks to its amenability to live imaging. Overall, due to the conserved features of OFT development among vertebrates (Grimes and Kirby, 2009), the zebrafish could help one to obtain new insights into the role of TGF-β signaling in OFT development and disease.

Here, we generated a zebrafish *alk5* mutant and observed a severe dilation of the developing OFT. We show that this phenotype results from early defects in EC proliferation and migration, followed by aberrant SMC proliferation and organization. Live imaging and transcriptomic analyses further reveal an Alk5-dependent alteration in extracellular matrix (ECM) composition. Notably, we show that restoring Alk5 in the endothelium is sufficient to rescue the OFT phenotype, including SMC organization defects. Moreover, we identify the ECM gene *fibulin-5* (*fbln5*) as a target of Alk5 signaling able to partially rescue the *alk5* mutant phenotype, providing a new therapeutic target for various aortic malformations.

## Results

### Lack of Alk5 causes specific defects in cardiac outflow tract formation

Mammalian *TgfbrI*/*Alk5* has two paralogs in zebrafish, *alk5a* and *alk5b*, and from multiple datasets (Pauli et al., 2012; Yang et al., 2013; Gauvrit et al., 2018; Mullapudi et al., 2018), we found *alk5b* to be the highest expressed paralog during embryogenesis. To investigate its expression, we performed *in situ* hybridization and generated a transgenic reporter line, *TgBAC*(*alk5b:eGFP*), by bacterial artificial chromosome (BAC) recombineering (Figure S1). We detected *alk5b* expression in the neural tube starting at 24 hours post fertilization (hpf) and in the gut at 72 hpf (Figure S1A-C’). Notably, in the developing cardiovascular system, *alk5b* reporter expression appeared to be restricted to the heart (Figure S1E, E’), as we could not detect GFP signal in any other vascular beds (Figure S1D, D’). Within the heart, *alk5b* becomes enriched in the outflow tract (OFT) (Figure 1A-B’’), where it is localized to both ECs and SMCs (Figure 1B’, B’’).

**Figure 1 –.**
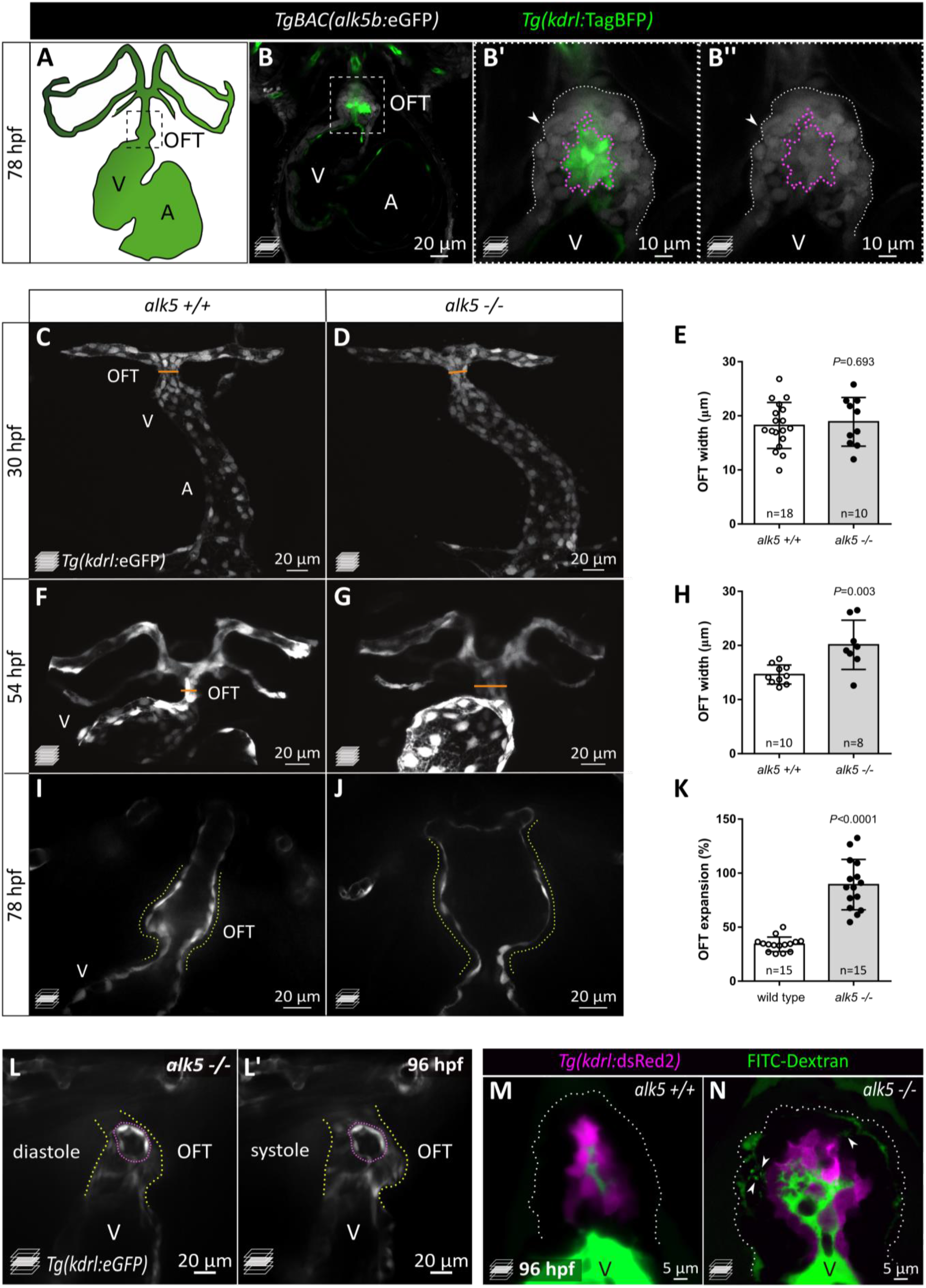
Loss of Alk5 function causes specific defects in cardiac outflow tract formation. A) Schematic of the zebrafish heart and connecting vessels at 78 hpf; ventral view; green, endothelium/endocardium. B-B’’) Confocal images showing *TgBAC*(*alk5b*:eGFP) (white) expression in the 78 hpf zebrafish heart. Green, ECs; arrowheads, SMCs; boxed area shown in B’ and B’’; white dotted line outlines the OFT; magenta dotted line outlines ECs. C-H) Confocal images (C, D, F, G) and quantification (E, H) of OFT width in wild-type and *alk5* mutant embryos at 30 (C-E) and 54 (F-H) hpf. Orange line shows OFT width quantified in E and H. I-K) Frames of confocal movies of beating hearts (I, J) and quantification (K) of OFT expansion at 78 hpf. Yellow dotted line outlines the OFT. L, L’) Frames of confocal movies of 96 hpf *alk5*-/- beating hearts during ventricular diastole (L) and systole (L’). Pink dotted lines outline EC ruptures. The OFT is outlined in yellow. C-L’) White, ECs. M, N) Confocal images of 96 hpf wild-type and *alk5* mutant OFTs, showing the accumulation (arrowheads) of FITC-Dextran (green) between the SMCs in mutant larvae. Magenta, ECs; dotted lines outline the OFT. C-G) Maximum intensity projections. B, I-N) Single confocal planes. E, H, K) Plot values represent mean ± SD; *P* values from *t*-tests. A-atrium, V-ventricle, OFT-outflow tract. Scale bars: B, C-L’) 20 μm; B’, B’’) 10 μm; M, N) 5 μm. See also Figure S1.

In order to investigate Alk5 function, we used CRISPR/Cas9 technology to generate mutants for *alk5a* and *alk5b*. We obtained a 4 bp and a 5 bp deletion in *alk5a* and *alk5b*, respectively, each leading to the predicted generation of truncated proteins, which lack the essential kinase domain (Figure S1F). Moreover, *alk5a* and *alk5b* mutant mRNA levels are decreased in the respective mutant fish while the other paralog does not appear to be upregulated, suggesting mutant mRNA degradation and a lack of transcriptional adaptation (El-Brolosy et al., 2019) (Figure S1G). Single *alk5a* and *alk5b* mutant larvae do not exhibit any gross morphological defects, other than the lack of inflation of the swim bladder in *alk5b* mutants (Figure S1H-J). Therefore, to achieve a complete blockade of Alk5 signaling, we generated *alk5a, alk5b* double mutants (*alk5a^-/-^;alk5b^-/-^*, hereafter referred to as *alk5* mutants). Loss of Alk5 function does not lead to early developmental defects until 72 hpf, when mutant larvae start exhibiting pericardial edema, evident at 96 hpf (Figure S1K), suggesting defective cardiac function. By analyzing heart morphology in live *Tg*(*kdrl:eGFP*) *alk5* mutant embryos, we observed a specific increase in OFT width by 54 hpf (Figure 1C-H). Live imaging on beating hearts showed that the dilation of the mutant OFT grows more severe with time, becoming more than twice as large as wild type by 78 hpf (+162%; Figure 1I-K; Video S1, S2). This expansion of the OFT is accompanied by its inability to pump blood into the connecting vessels, leading to retrograde flow into the ventricle (Video S3).

In 78 hpf wild-type zebrafish, the OFT is connected to the aortic arches by a single vessel, the ventral aorta (VA) (Figure S1L). *alk5* mutants fail to form this vessel, leading to two independent connections from the OFT to the left and right aortic arches (Figure S1M). Furthermore, we occasionally observed in mutant OFTs ruptures in the endothelial layer (Figure 1L, L’; Video S4), likely resulting from a leaky endothelium. Indeed, 10 minutes after dextran injection into the circulation, 7 out of 11 mutant larvae displayed dextran accumulation between the ECs and SMCs and in the interstitial space amongst SMCs, a phenotype not observed in wild types (n=10; Figure 1M, N). Remarkably, the cardiovascular defects in *alk5* mutant fish are restricted to the OFT and the VA, while all other vascular beds appear morphologically unaffected. The diameter of the dorsal aorta (DA) appears unaffected in *alk5* mutants compared with wild-type siblings at 56 and 96 hpf (Figure S1N), and the atrium, ventricle and atrioventricular valve appear correctly shaped at 78 hpf (Figure S1O, P).

Taken together, these results identify a previously unknown and specific requirement for Alk5 in OFT morphogenesis, structural integrity, and functionality.

### Alk5 restricts EC proliferation in the cardiac outflow tract and promotes EC migration towards the ventral aorta

Given the increased size of the mutant OFT, we asked whether this phenotype was accompanied by an increase in cell number. In wild-type animals, the average number of OFT ECs increases from 21, at 36 hpf, to 45, at 72 hpf (Figure 2A, B). Consistent with the absence of a morphological phenotype, the mutant OFT, at 36 hpf, was composed of an average of 22 ECs, similar to wild type (Figure 2B). However, the number of ECs in mutant OFT diverges substantially over time, and by 72 hpf twice as many ECs were observed in mutants compared to wild types (85 ECs; Figure 2B). To investigate the underlying cause of this increase, we performed EdU labeling to assess EC proliferation. In the mutant OFTs, we found that, by 36 hpf, ECs were already more likely to undergo cell cycle reentry than ECs in wild-type OFTs (Figure 2C-E). This abnormal increase in the number of EdU^+^ ECs in mutant OFTs becomes even more pronounced at later stages (48-72 hpf; Figure 2F-H).

**Figure 2 –.**
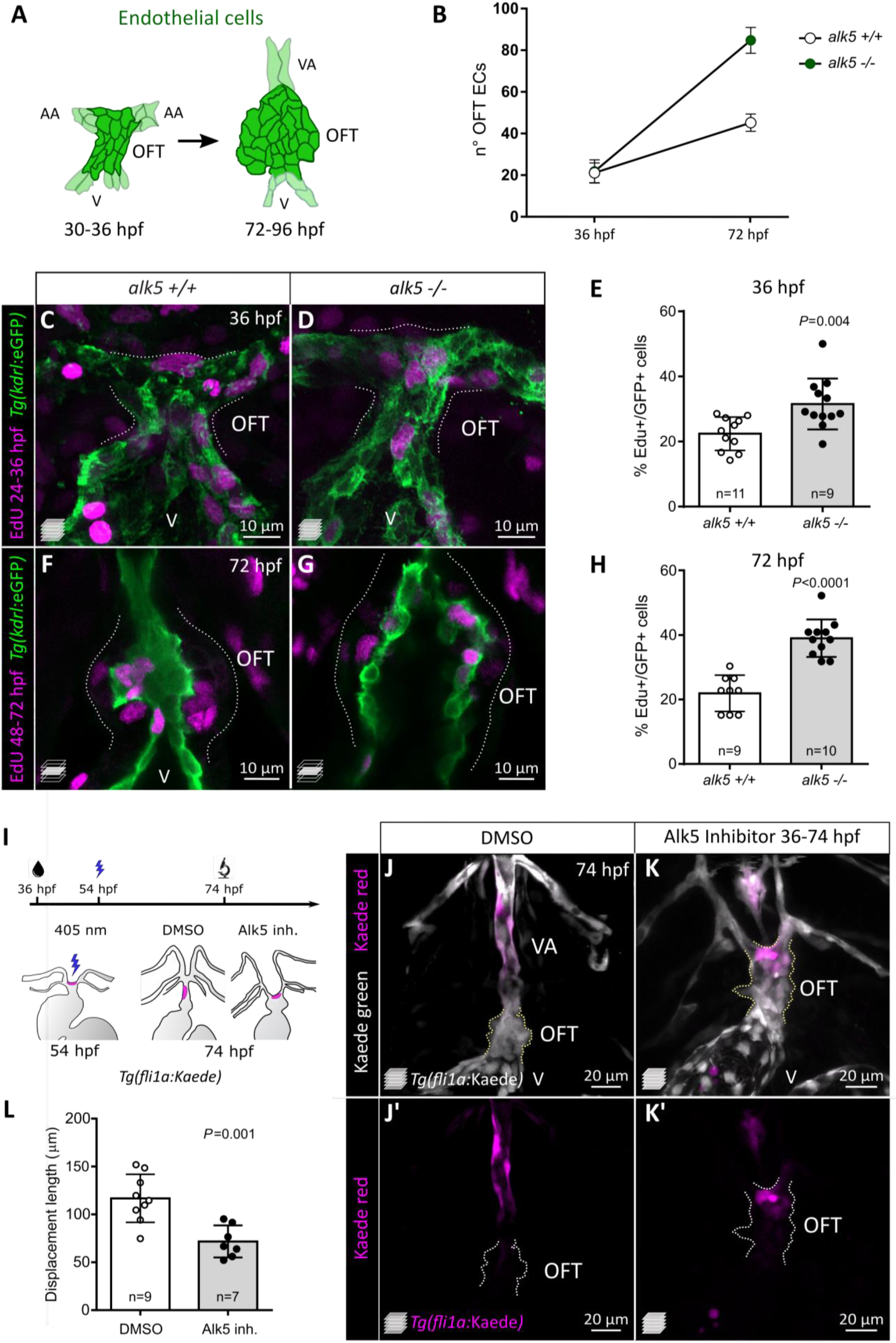
Alk5 restricts EC proliferation in the cardiac outflow tract and promotes EC migration towards the ventral aorta. A) Schematics of OFT ECs at 36 and 72 hpf. B) Quantification of EC number (darker cells shown in A) in 36 and 72 hpf wild-type and *alk5* mutant OFTs (36 hpf: n=11; 72 hpf: n=6). C-H) Confocal images (C, D, F, G) and quantification (E, H) of the percentage of EdU^+^ ECs in wild-type and *alk5* mutant *Tg(kdrl:eGFP)* animals. Dotted lines outline the OFT. I) Protocol used for photoconversion experiment. J-K’) Confocal images of the OFT in 74 hpf *Tg(fli1a:Kaede)* larvae treated with DMSO or Alk5 inhibitor. Magenta, photoconverted ECs; dotted lines outline the OFT. L) Quantification of the distance covered by photoconverted ECs between 54 and 72 hpf in control and Alk5 inhibitor treated larvae. C, D, J-K’) Maximum intensity projections. F, G) Single confocal planes. B, E, H, L) Plot values represent means ± SD; *P* values from *t*-tests. VA-ventral artery, OFT-outflow tract, V-ventricle. Scale bars: C-G) 10 μm; J, K) 20 μm. See also Figure S2.

Along with proliferation, EC migration has been implicated in the patterning and formation of blood vessels, such as in the case of the aortic arches (Rochon et al., 2016). Therefore, we set to investigate if the absence of the VA in *alk5* mutants could be attributed to a defect in EC migration. To assess EC migration in the absence of Alk5 activity, we performed photoconversion of OFT ECs in control and Alk5 inhibitor-treated larvae. Treatment with 2.5 μM Alk5 inhibitor, starting at 36 hpf (Figure S2A), was found to cause increased OFT width at 54 and 78 hpf (Figure S2B-F) and VA patterning defects (Figure S2D, E), thus phenocopying *alk5* mutants. Using the Alk5 inhibitor with the *Tg*(*fli1a:Kaede*) transgenic line, we aimed to track the migration of photoconverted ECs in the region of the OFT. Since the population of ECs which gives rise to the VA was not previously characterized, we photoconverted ECs at 54 hpf in different regions of the OFT: the proximal region close to the bulbo-ventricular (BV) valve (Figure S2G, H), or the most distal region between the aortic arches (Figure S2J, K). In wild-type fish, proximal ECs remained in the OFT close to the photoconversion site up to 74 hpf (Figure S2H, I), whereas distal ECs were invariably found to move into the VA (Figure S2J; Figure 2I-J’). In contrast, upon Alk5 inhibition, also distal ECs remained in the OFT (Figure S2L; Figure 2K, K’). By 74 hpf, these ECs were displaced to a lesser extent compared to control larvae (Figure 2L). To observe EC migration at high resolution, we recorded time-lapse movies from 56 to 74 hpf following photoconversion (Video S5, S6). While photoconverted cells in control larvae extended rostrally to form the VA (Video S5), ECs with reduced Alk5 activity did not appear to move from their original position in the OFT, resulting in the lack of VA formation (Video S6).

Altogether, these data indicate that Alk5 plays a role in restricting EC proliferation in the OFT and promotes EC migration to form the ventral artery.

### Alk5 promotes the formation and stability of the outflow tract wall, regulating SMC proliferation and organization

During early larval stages, the OFT becomes covered by SMCs, which allow it to buffer the high blood pressure caused by ventricular contractions. In order to visualize SMCs, we used the *Tg*(*pdgfrb:eGFP*) line, which labels these cells before they differentiate into more mature SMCs and start expressing established markers such as Acta2 (smooth muscle actin; Figure 3A-B’). We observed that the OFT endothelium in 75 hpf wild-type larvae is surrounded by an average of 90 ± 2 *pdgfrb*^+^ cells, organized in 2 to 3 compact layers with limited extracellular space (Figure 3A, A’, C). In contrast, in *alk5* mutants we observed a reduced number of *pdgfrb*^+^ SMCs (65 ± 2, −27%), wider extracellular space, and disorganized cell layers around the OFT (Figure 3B-C). The reduction in the total number of SMCs is likely caused by a proliferation defect, as confirmed by EdU incorporation experiments performed at early larval stages (−53%; 48-72 hpf, Figure 3D). Furthermore, using TUNEL staining (data not shown), we excluded SMC death by apoptosis.

**Figure 3 –.**
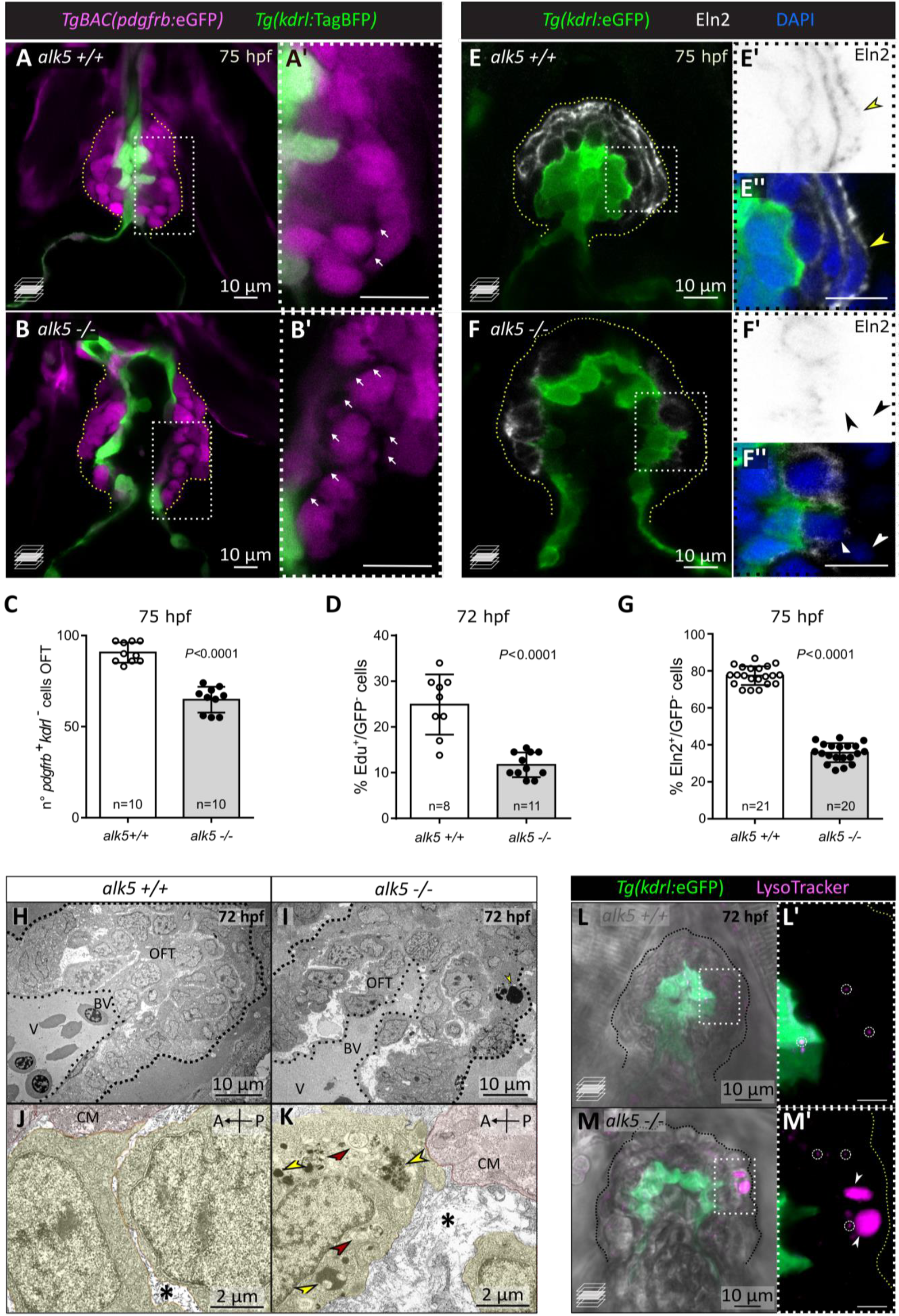
Alk5 regulates SMC and ECM organization in the OFT. A-C) Confocal images (A-B’) and quantification (C) of SMCs in 75 hpf wild-type and *alk5* mutant larvae. Magenta, SMCs; white arrows, extracellular space between SMCs; boxed areas shown in A’ and B’; dotted lines outline the OFT. D) Percentage of EdU^+^ SMCs in 72 hpf wild-type and *alk5* mutant larvae. E-F’’) Confocal images of wild-type and *alk5* mutant larvae immunostained for Elastin2 (Eln2) at 75 hpf. White arrowheads, SMCs devoid of Eln2; yellow arrowheads SMCs surrounded by Eln2 immunostaining; boxed areas shown in E’, E’’, F’ and F’’; images in E’ and F’ are shown with inverted colors; dotted lines outline the OFT. G) Quantification of the percentage of SMCs surrounded by Eln2 immunostaining (per sagittal plane) at 75 hpf. H-K) TEM images of wild-type and mutant OFTs at 72 hpf at different magnifications. Yellow, SMCs; red, cardiomyocytes close to the BV canal; asterisks, extracellular space; arrows, electron-dense (red) and double-membraned (yellow) vacuoles; dotted lines outline the OFT. L-M’) Confocal images of wild-type and *alk5* mutant animals treated with LysoTracker, labelling lysosomes (small, circles; big, arrowheads); boxed areas are shown in L’ and M’; dotted lines outline the OFT. C, D, G) Plot values represent means ± SD; *P* values from *t*-tests. A-anterior, P-posterior, BV-bulbo-ventricular canal, CM-cardiomyocyte. Scale bars: A-I, L-M’) 10 μm; J, K) 2 μm.

In order to form a compact yet elastic wall, the SMCs are embedded within a specialized extracellular matrix (ECM), which provides essential biomechanical support as well as signaling cues to the SMCs (Raines, 2000). To assess whether and how ECM structure and composition were affected in *alk5* mutants, we analyzed the localization of Elastin2 (Eln2; Figure 3E-G), a major ECM component in the OFT (Miao et al., 2007). We observed that in wild-type larvae 77.9% of SMCs were surrounded by a continuous layer of Eln2, while in *alk5* mutants only 33.1% were surrounded (Figure 3G). Moreover, in *alk5* mutant OFTs Eln2 localizes in small disrupted clusters, compared to the bundle localization characteristic of elastic fibers in wild types (Figure 3E-F’’).

Next, we used transmission electron microscopy (TEM) to investigate OFT ultrastructure (Figure 3H-K). Even in a low magnification view of the mutant OFTs, we could observe a greatly widened extracellular space surrounding the SMCs (Figure 3I). At higher resolution, we observed that the ECM in wild-type larvae consisted mainly of thin layers (Figure 3J), while in *alk5* mutants it consisted of broader electron-negative spaces (Figure 3K). Moreover, SMCs in the external layer of the mutant OFTs exhibited cytoplasmic inclusions (Figure 3K), such as electron-dense lysosomes (as confirmed by LysoTracker staining; Figure 3L-M’), and double-membraned vacuoles, resembling autophagosomes.

Overall, the loss of Alk5 function results in the formation of a defective SMC wall, supported by a structurally impaired ECM.

### Endothelial Alk5 is sufficient to restore cardiac outflow tract wall formation and function

The earliest phenotype in *alk5* mutants is observed during the EC proliferation and migration stages, thereby preceding the formation of the SMC wall. Therefore, to investigate Alk5 requirement in the endothelium, we generated a transgenic line overexpressing *alk5b* specifically in ECs. We used the *fli1a* promoter to drive *alk5b* expression (Figure 4A) and validated the specificity of *Tg(fli1a:alk5b-mScarlet)^bns421^* expression in endothelial and endocardial cells (Figure S3A-A’’). We found that fish overexpressing *alk5b* in wild-type ECs were healthy, fertile and did not display obvious cardiovascular phenotypes (Figure S3A). Notably, when *alk5b* was overexpressed in the ECs in a mutant background, it was sufficient to efficiently restore cardiac function, as indicated by the lack of pericardial edema (Figure 4B, C). Indeed, *alk5* mutants carrying the rescue transgene exhibited a wild-type morphology and function of the OFT and connecting vessels (Figure 4D-F), including a single VA (Figure S3B-D) and a wild-type like OFT expansion (Figure 4G; Video S7). Occasionally, the restored cardiac function and blood flow allowed a few (10.2%, n=108) of the *alk5* rescued mutants to inflate their swim bladder and survive until 9 days post fertilization (dpf) (Figure S3E, F). However, despite the vascular rescue, the fish do not survive to adulthood, presumably due to the lack of Alk5 signaling in other tissues, as suggested by an altered morphology of the head (Figure S3F).

**Figure 4 –.**
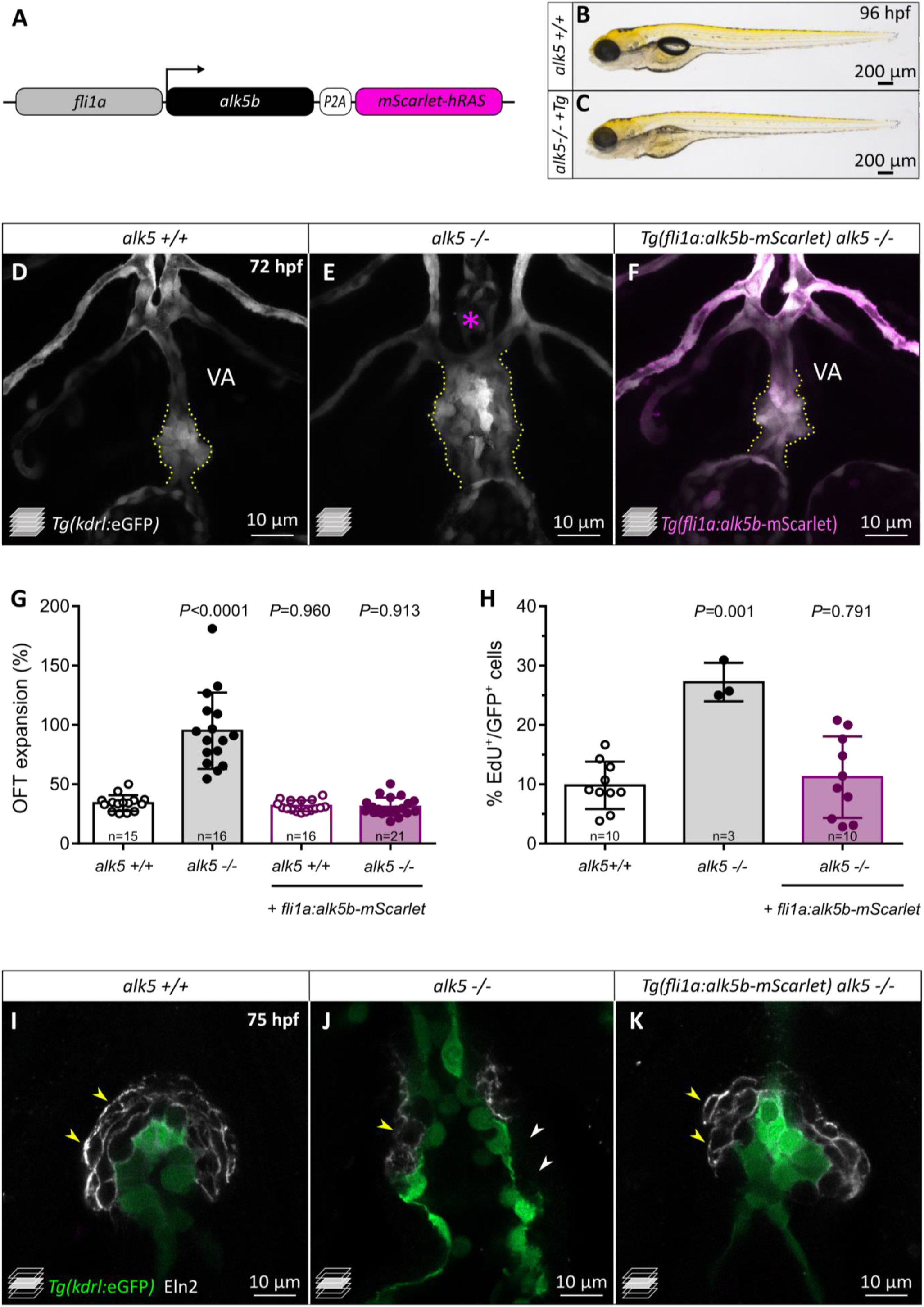
Endothelial Alk5 is sufficient to restore OFT wall formation and function. A) Schematic of the construct used for endothelial-specific rescue experiments. B, C) Brightfield images of 96 hpf wild-type and *alk5* mutant larvae carrying the EC-specific rescue transgene (*Tg*). D-F) Confocal images of the OFT in 72 hpf *Tg(kdrl:eGFP)* larvae, showing the morphological rescue in *Tg(fli1:alk5b-mScarlet) alk5* mutants (F). Asterisk indicates the absence of the VA in *alk5* mutants; dotted lines outline the OFT. G) Percentage of OFT expansion in 78 hpf wild types, *alk5* mutants, and *Tg(fli1:alk5b-mScarlet) alk5* mutants. H) Percentage of EdU^+^ ECs in 72 hpf wild types, *alk5* mutants, and *Tg(fli1:alk5b-mScarlet) alk5* mutants. I-K) Confocal images of 75 hpf *Tg(kdrl:eGFP)* larvae immunostained for Eln2. Arrowheads, SMCs devoid of (white) or surrounded by (yellow) Eln2 immunostaining. G, H) Plot values represent means ± SD; *P* values from One way ANOVA test. VA, ventral artery; *Tg*, *Tg(fli1:alk5b-mScarlet)*. Scale bars: B, C) 200 μm; D-F, I-K) 10 μm. See also Figure S3.

In order to characterize the EC-specific OFT rescue at a cellular level, we performed EdU incorporation experiments and observed that the number of proliferating ECs was reduced in transgenic *alk5* mutant larvae compared to *alk5* mutants not carrying the transgene (Figure 4H). Remarkably, the overexpression of *alk5b* in the endothelium of *alk5* mutant larvae also restored the formation of the SMC wall (Figure 4I-K). In fact, 75 hpf transgenic larvae exhibited SMCs organized in multiple layers around the OFT (Figure 4K), and 70.8% (n=16) of these cells were surrounded by uniform Eln2 immunostaining (Figure S3G), a percentage similar to that observed in wild types (77,6%, n=12).

Overall, these data suggest that endothelial Alk5 signaling is sufficient to restore OFT morphology and function, including SMC wall formation.

### Alk5 signaling regulates ECM gene expression

We hypothesized that Alk5 is required in the OFT endothelium and that it controls an expression program modulating the structural integrity of the OFT. To identify candidate effector genes, we performed a transcriptomic analysis using manually extracted embryonic hearts, including the OFTs, at 56 hpf in control and Alk5 inhibitor treated embryos (Figure 5A). We chose 56 hpf as the developmental stage for this analysis in order to avoid secondary effects deriving from OFT malfunction.

**Figure 5 –.**
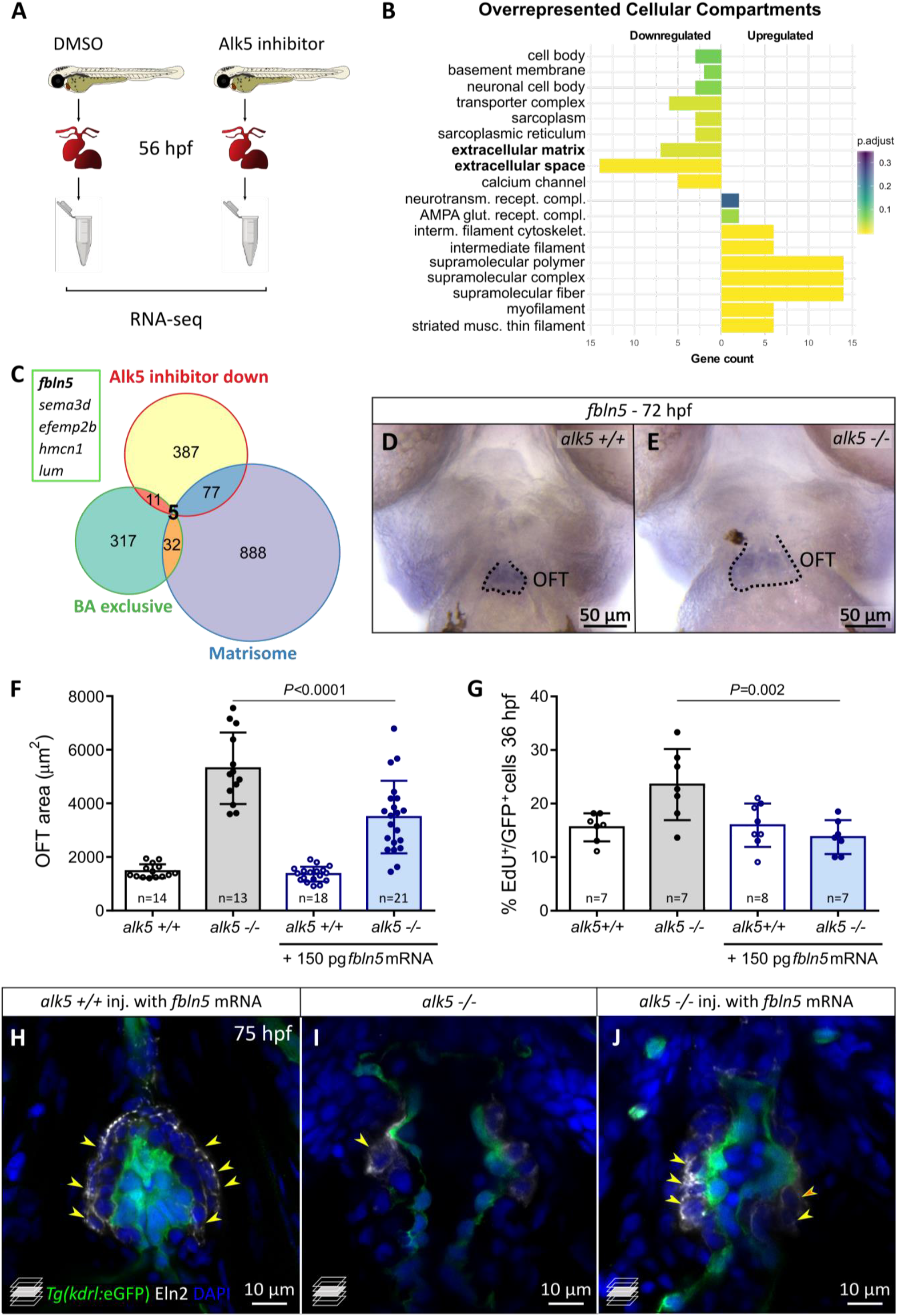
Alk5 signaling regulates ECM component genes of the outflow tract, including *fbln5*. A) Schematic showing the setting of the RNA-seq on extracted embryonic hearts (56 hpf). B) Gene ontology bar plot, showing the most overrepresented cellular compartments up- or down-regulated in the inhibitor treated embryos compared to controls. C) Venn diagram depicting the genes downregulated (log2FC < −0.585) in Alk5 inhibitor treated embryonic hearts, exclusively expressed in the adult zebrafish BA (Singh et al., 2016), and part of the zebrafish matrisome (Nauroy et al., 2018). The 5 genes in the intersection are listed in the box. D, E) Whole-mount *in situ* hybridization of *fbln5* expression in the OFT of 72 hpf wild-type and *alk5* mutant larvae. Dotted lines outline the OFT. F) Quantification of the maximum OFT area during ventricular systole in 78 hpf *alk5* wild-type and mutant larvae following injection of *fbln5* mRNA. G) Percentage of EdU^+^ ECs in 36 hpf wild-type and *alk5* mutant *Tg(kdrl:eGFP)* embryos following injection of *fbln5* mRNA. H-J) Confocal images of 75 hpf larvae immunostained for Eln2, following injection of *fbln5*. Yellow arrowheads, SMCs surrounded by Eln2 immunostaining. OFT, outflow tract; BA, bulbus arteriosus. F, G) Plot values represent means ± SD; *P* values from One way ANOVA test. Scale bars: D, E) 50 μm; H-J) 10 μm. See also Figure S4.

Analysis of the Alk5 inhibitor treated samples revealed 955 differentially expressed genes (DEGs), 480 of which were downregulated compared to control (Figure S4A-C). Gene ontology analysis showed enrichment of genes implicated in the response to TGF-β signaling among the downregulated genes (Figure S4B), supporting the specificity of the inhibitor treatment. Notably, genes downregulated upon Alk5 inhibition include multiple ECM components and the related Gene ontology categories are amongst the most enriched (Figure 5B). In order to identify extracellular proteins that might function in the signaling between ECs and SMCs, we compared the list of downregulated genes with the secreted factor genes from the zebrafish matrisome (Nauroy et al., 2018) (Figure 5C). Among the 82 genes identified, 5 of them were specifically expressed in the OFT in adult zebrafish (Singh et al., 2016), with *fbln5* being the most downregulated one upon Alk5 inhibition (Figure 5C, S4C). Fbln5 is an ECM protein which mediates cell-to-matrix adhesion and has several functions, including the inhibition of EC proliferation *in vitro* and the organization of the elastic lamina (Sullivan et al., 2007; Yanagisawa et al., 2009). Interestingly, mouse *Fbln5* is expressed in both fibroblasts/SMCs and ECs (Figure S4D) (Tabula Muris et al., 2018), also specifically in the embryonic OFT and aorta (Liu et al., 2019). Thus, we first confirmed that *fbln5* mRNA levels were also decreased in zebrafish *alk5* mutant hearts by RT-qPCR (Figure S4E) and, next, we observed that *fbln5* expression was indeed restricted to the OFT in both wild-type and *alk5* mutant larvae at 72 hpf (Figure 5D, E).

*fbln5* specific expression in the developing OFT and its downregulation in *alk5* mutants suggest a potential role downstream of TGF-β signaling during OFT formation. Therefore, we injected *fbln5* mRNA in *alk5* mutants and assessed the size of the OFT during ventricular systole at 78 hpf (Figure S4F). Despite the transient effect of injected mRNA, the injection of *fbln5* mRNA partially rescued the size of the OFT in 15 out of 21 *alk5* mutants, while it did not have any effect in wild-type siblings (Figure 5F). Moreover, *alk5* mutants injected with *fbln5* mRNA exhibited a percentage of EdU^+^ ECs in early embryonic stages (16,0%; 24-36 hpf, Figure 5G), which is comparable with wild types (15,6%). Additionally, *fbln5* mRNA injections into *alk5* mutants led to a more organized Eln2 immunostaining around the OFT SMCs compared to non-injected mutants (Figure 5H-J; S4G-I).

Overall, these data suggest a potential role for Fbln5 downstream of Alk5 signaling in OFT morphogenesis, by modulating EC proliferation and elastin organization.

## Discussion

Taking advantage of the zebrafish model, we report an important role for the TGF-β receptor I Alk5 in the morphogenesis of the OFT. We show that TGF-β signaling through Alk5 restricts EC proliferation and promotes EC migration towards the ventral artery during early embryonic stages. Furthermore, loss of Alk5 causes defects in the SMC wall surrounding the OFT. The combination of these phenotypes leads to defects in OFT structural integrity, eventually resulting in vessel dissection, reminiscent of human pathologies. Notably, restoring Alk5 expression in the endothelium is sufficient to rescue the OFT defects, suggesting a critical role for Alk5 in ECs.

### Cell-specific role of Alk5 signaling in cardiac outflow tract development

The TGF-β pathway’s multi-functional role results in many context- and tissue-dependent functions, which are difficult to unravel (Pardali et al., 2010). Most studies on diseases of the great arteries have aimed to understand the role of TGF-β in SMCs (Li et al., 2014; Yang et al., 2016; Zhang et al., 2016; Perrucci et al., 2017; Takeda et al., 2018), overlooking its function in the endothelium. In EC biology, Alk5 function has been mostly studied *in vitro*, where it has been shown to maintain EC quiescence (Goumans et al., 2002; Goumans et al., 2003; Lebrin et al., 2004; van Meeteren and ten Dijke, 2012; Maring et al., 2016). In the zebrafish model, we confirmed a pivotal role for Alk5 in restricting EC proliferation *in vivo*. We found that ECs lacking Alk5 function also exhibited impaired migration, possibly due to the downregulation of *alk1* (another type I TGF-β receptor gene) and *sema3d*, which have been previously implicated in EC migration (Hamm et al., 2016; Rochon et al., 2016). Moreover, time-lapse imaging combined with photoconversion experiments identified a selected population of OFT ECs giving rise to the ventral aorta via a migration process, which is impaired in *alk5* mutants.

Importantly, the early EC defects together with the endothelial-specific rescue data suggest that ECs are primarily responsible for the OFT phenotype. Remarkably, while in mouse, *Alk5* expression appears to be enriched in the aortic SMCs (Seki et al., 2006), a few studies have suggested a pivotal role for TGF-β in ECs (Sridurongrit et al., 2008; Bochenek et al., 2020). For example, although *Tgfbr2* deletion in SMCs leads to OFT expansion in mouse (Jaffe et al., 2012), only EC-specific *Alk5* KO mice recapitulate the early cardiovascular defects of mice carrying a global *Alk5* mutation (Carvalho et al., 2007; Sridurongrit et al., 2008; Dave et al., 2018). This discrepancy indicates distinct requirement for various TGF-β receptors in different cell types and developmental stages.

Notably, in *alk5* mutant zebrafish larvae, the SMCs appears after the EC phenotype. In stark contrast to the effect in ECs, SMCs in *alk5* mutants exhibit reduced proliferation and therefore reduced coverage of the OFT, explaining the lost integrity of the wall. This reduced SMC coverage is in agreement with the vascular defects reported in mice lacking Alk5 signaling globally, as well as in patients affected by aortic aneurysms (Larsson et al., 2001; Jana et al., 2019).

### Endothelium-smooth muscle interplay in the cardiac outflow tract and aorta

Surprisingly, we found that *alk5* overexpression in ECs was sufficient to restore SMC wall formation. Although these results do not resolve whether ECs are the only cell type in which Alk5 is necessary for OFT morphogenesis, they suggest that the SMC phenotype could be a secondary effect from the crucial cross-talk between these two cell types (Gaengel et al., 2009; Lilly, 2014; Stratman et al., 2017; Perbellini et al., 2018). Supporting the importance of this cross-talk, it has recently been described that pericyte-specific *Alk5* deletion causes an enlargement of brain capillaries in mice. In particular, ALK5 in pericytes interferes with ECM degradation processes, leading to an EC hyperproliferation state (Dave et al., 2018). Moreover, defective interactions between ECs and SMCs have been implicated in different pathologies including pulmonary hypertension, atherosclerosis and arteriovenous malformations (e.g., hereditary hemorrhagic telangiectasia) (Mancini et al., 2009; Gao et al., 2016; Cunha et al., 2017; Li et al., 2018).

The communication between ECs and SMCs can be carried out in different ways, such as physical contact, exchange of signaling cues and ECM deposition (Gaengel et al., 2009; Lilly, 2014; Li et al., 2018; Sweeney and Foldes, 2018), all of which are likely to play a role in *alk5* mutant phenotype. In addition, during development, SMCs are recruited to the vessels once the initial establishment of the endothelial layer is completed (Stratman et al., 2017; Sweeney and Foldes, 2018). Thus, one can speculate that enhanced EC proliferation and remodeling in *alk5* mutants could represent a less mature state of the vessel, thus inhibiting SMC coverage of the OFT. Later on, a reduced SMCs coverage might sustain the EC hyperproliferation in a feedback loop contributing to the severity of the phenotype.

Overall, we suggest that the severe SMC phenotype directly causing the functional OFT defects and leading to aortic aneurysms in patients could have masked the important role of the endothelium in the etiology of these pathologies. Therefore, it will be important to further characterize the EC role in SMC stabilization and vascular wall integrity.

### Identifying new molecular regulators of cardiac outflow tract development and disease

Analyses at several different cellular and molecular levels provided insights into the structural changes directly linked to the functional phenotype in *alk5* mutants. Together with the observed defective elastic lamina and altered intercellular space, the transcriptomic data identified differential expression of several ECM component genes following manipulation of Alk5 function. The ECM is secreted by both ECs and SMCs and is a source of signaling mediators crucial for their interaction (Kelleher et al., 2004; Davis and Senger, 2005). One of the most promising ECM genes downregulated after Alk5 inhibition was *fbln5*, encoding an integrin-binding extracellular protein (Nakamura et al., 1999). *Fbln5* is expressed, and secreted, by both ECs and fibroblasts or SMCs (Nakamura et al., 1999; Tabula Muris et al., 2018; Liu et al., 2019) and, in zebrafish, it serves as a very specific marker for the OFT (Figure 5D). FBLN5 has multiple reported functions, as assembling the elastic lamina surrounding SMCs (Nakamura et al., 2002; Chapman et al., 2010) and promoting EC-to-ECM attachment *in vitro* (Preis et al., 2006; Williamson et al., 2007). The adhesion of ECs to the matrix mediated by FBLN5 appears essential to restrict their proliferation, suggesting FBLN5 as an anti-angiogenic factor (Albig and Schiemann, 2004; Sullivan et al., 2007). Moreover, the deregulation of *FBLN5* has been associated with abdominal aortic aneurysms, but the underlying mechanism is still unclear (Orriols et al., 2016). By enhancing the levels of *fbln5* in *alk5* mutants, we were able to partially restore the OFT expansion and the Eln2 coverage of SMCs. Notably, the injection of *fbln5* mRNA also rescued the EC hyperproliferation phenotype of *alk5* mutants at a stage preceding the formation of the SMC wall. These data are consistent with FBLN5 reported role in restricting EC proliferation, likely acting via cell-to-matrix adhesion mechanisms. While we have not identified the exact cell type expressing *fbln5*, it is important to note that multiple cells secrete ECM components during OFT morphogenesis. Nevertheless, it is conceivable that ECs in the OFT could initiate the generation of a local extracellular environment, the control of which is then taken over by SMCs.

Given the prevalence of OFT-related cardiovascular diseases, there is a need for multiple model systems that allow one to investigate the causes of these pathologies at the cellular and molecular levels, and the zebrafish could complement the mouse in this regard. Furthermore, the overlap of our transcriptomic data with the genes associated with aortic aneurysm (Brownstein et al., 2017; Kim and Stansfield, 2017), and the similarities between the endothelial ruptures in *alk5* mutants and those of human aortic dissection suggest that the zebrafish could serve as a valuable model for aneurysm research.

## Acknowledgments

We would like to thank Radhan Ramadass for critical help with microscopy and all the fish facility staff for technical support; Rashmi Priya for the *myl7:mScarlet-Hsa.HRAS* plasmid and valuable suggestions; Matteo Perino for help with the analyses, discussions and critical comments on the manuscript; Sri Teja Mullapudi, Paolo Panza, Josephine Gollin for suggestions and critical comments on the manuscript. Research in the Stainier lab is supported in part by the Max Planck Society, DFG (Sonderforschungsbereich (SFB 834) and the European Union (ERC).

## Author contributions

Conceptualization, G.L.M.B, C.S.M.H, A.B.B. and D.Y.R.S.; Methodology, G.L.M.B, J.P., S.G.; Validation, G.L.M.B.; Formal Analysis, G.L.M.B., S.G.; Investigation, G.L.M.B., C.S.H.M., J.P., S.G.; Writing – Original Draft, G.L.M.B. and D.Y.R.S.; Writing – Reviewing & Editing, all; Visualization, G.L.M.B.; Supervision, C.S.M.H and D.Y.R.S; Project Administration, D.Y.R.S.; Funding Acquisition, D.Y.R.S.

---

**Figure S1 –.**
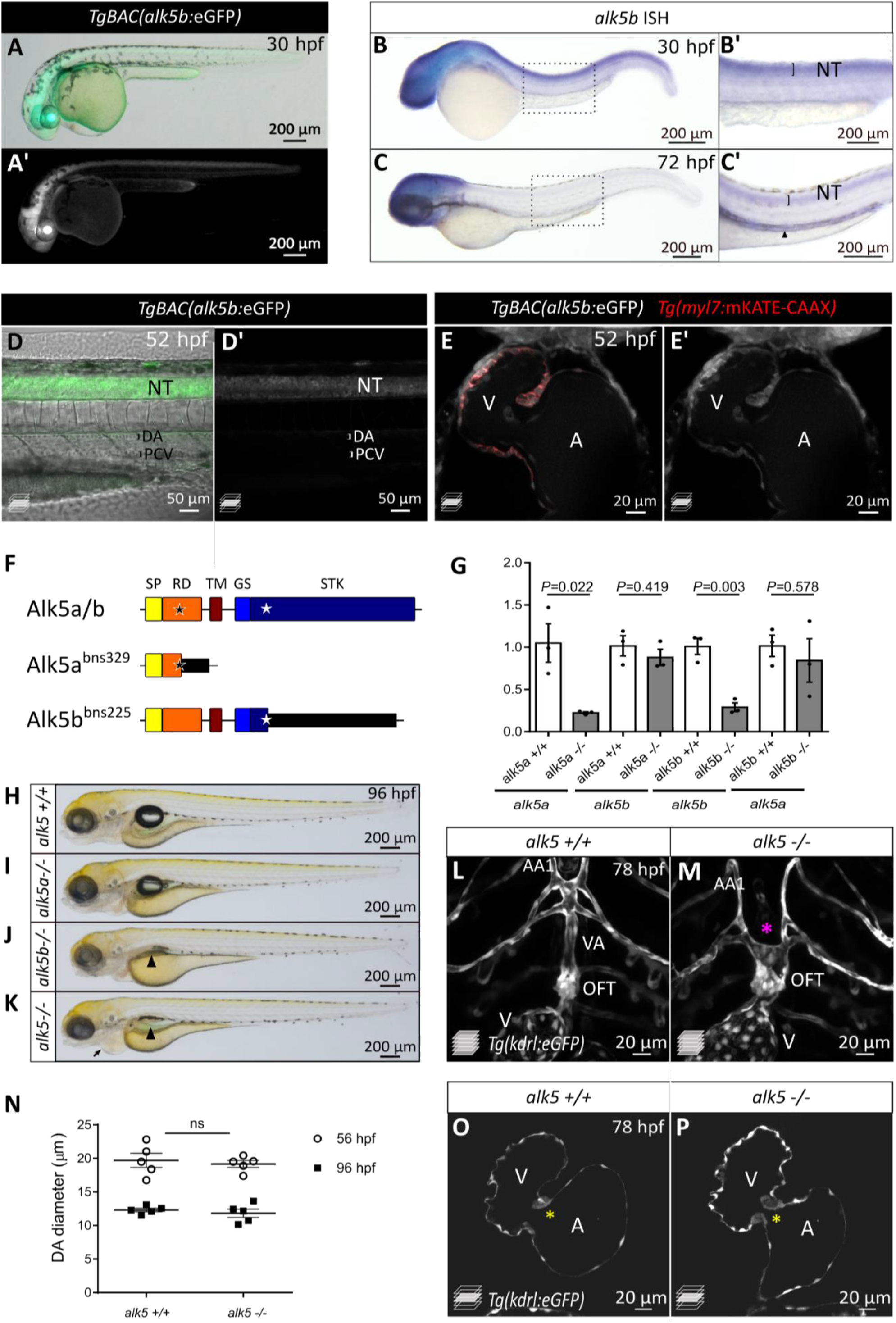
Alk5 expression and function in zebrafish embryos and larvae – related to Figure 1. A-A’, D, D’, E, E’) Confocal images showing *TgBAC(alk5b:eGFP)* (white) expression in 30 (A) and 52 (D-E’) hpf zebrafish embryos. The expression is localized in the neural tube (D) and the heart (E). B, C) *In situ* hybridization showing the expression of *alk5b* in 30 and 72 hpf animals. Arrowhead, gut; boxed area shown in B’ and C’. F) Schematics of wild-type and mutated Alk5 proteins. Black and white stars indicate the mutations in *alk5a* and *alk5b*, respectively. G) *alk5a* and *alk5b* mRNA levels in 30 hpf wild-type, *alk5a* and *alk5b* single mutants; means ± SD; *P* values from *t*-tests. H-K) Brightfield images of 96 hpf wild-type and mutant larvae. Arrowhead, lack of swim bladder; arrow, pericardial edema. L, M) Confocal images of OFT and connecting vessels in 78 hpf wild-type and mutant larvae. Asterisk, absence of the VA in *alk5* mutants. N) Quantification of DA diameter in 56 and 96 hpf wild-type and *alk5* mutant animals. The same animals were analyzed at the two different time-points; means ± SD; *P* values from *t*-tests. O, P) Confocal images of 78 hpf wild-type and mutant hearts. Asterisk, AV valve. SP-signaling peptide, RD-receptor domain, TM-transmembrane domain, GS-glycine-serine rich domain, STK-serine-threonine kinase domain, A-atrium, V-ventricle, NT-neural tube, DA-dorsal aorta, PCV-posterior cardinal vein, OFT-outflow tract, VA-ventral artery, AA1-aortic arch 1. Scale bars: A-C’, H-K) 200 μm; D, D’) 50 μm; E, E’, L-P) 20 μm.

**Figure S2 –.**
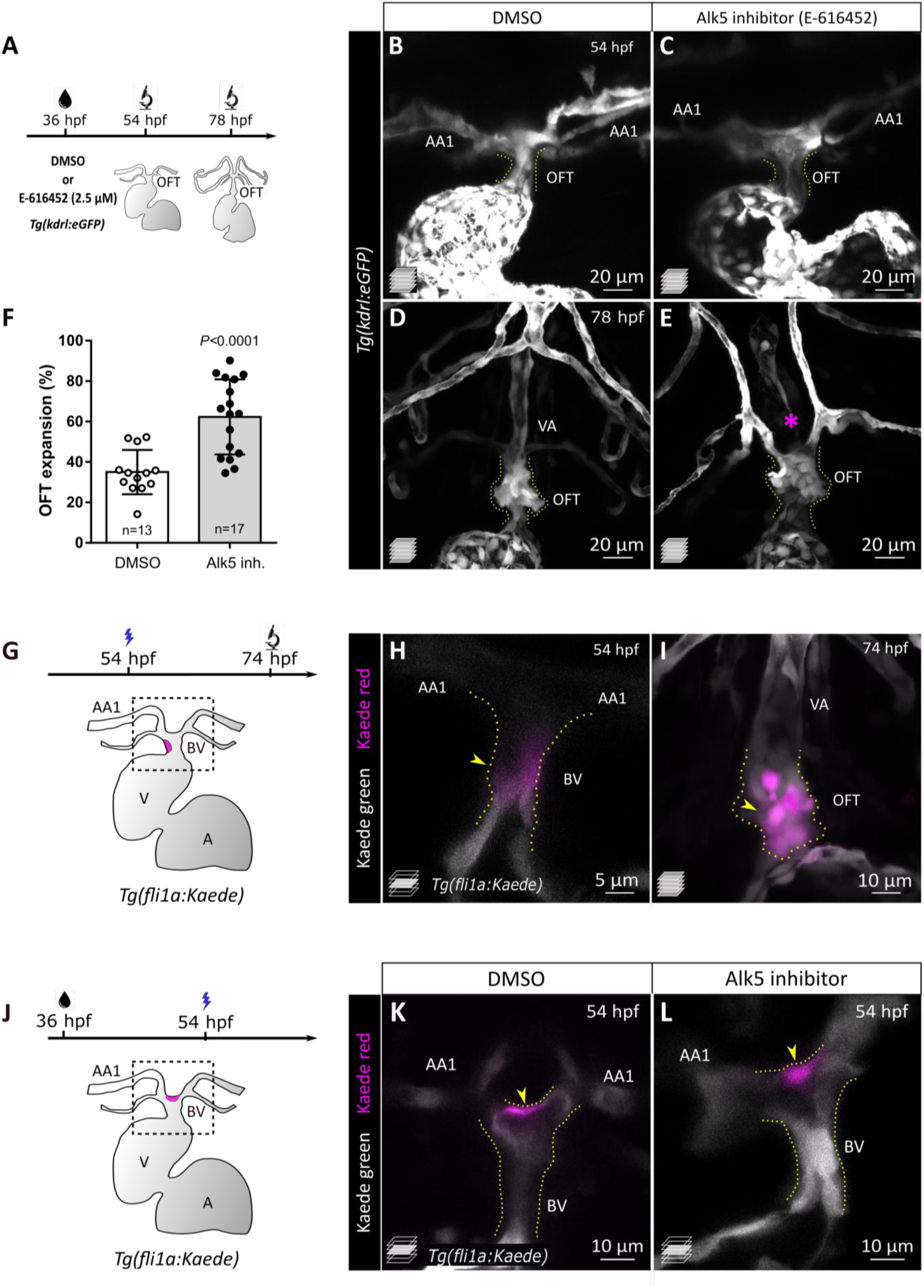
Phenocopy of *alk5* mutants by Alk5 inhibitor treatment – related to Figure 2. A) Protocol used for Alk5 inhibitor treatment. B-E) Confocal images of *Tg(kdrl:eGFP)* animals treated with DMSO or Alk5 inhibitor (E-616452) at 36 hpf and analyzed at 54 (B, C) or 78 (D, E) hpf. Asterisk, absence of the VA in Alk5 inhibitor treated larvae. F) Percentage of OFT expansion in 78 hpf control and Alk5 inhibitor treated larvae; means ± SD; *P* values from *t*-test. G) Schematics of the area photoconverted in H. H, I) Confocal images of untreated photoconverted *Tg(fli1a:Kaede)* embryos at 54 (H) and 74 (I) hpf. J) Schematics of the area photoconverted in K, L. K, L) Confocal images of 54 hpf photoconverted *Tg(fli1a:Kaede)* control or Alk5 inhibitor treated embryos. H-L) Magenta, photoconverted ECs in the OFT (yellow arrowheads). B-L) Dotted lines outline the OFT. H, K, L) Single confocal planes. B-E, I) Maximum intensity projections. AA1-1° aortic arch, VA-ventral artery, OFT-outflow tract, BV-bulbo-ventricular canal. Scale bars: B-E) 20 μm; I-L) 10 μm; H) 5 μm.

**Figure S3 –.**
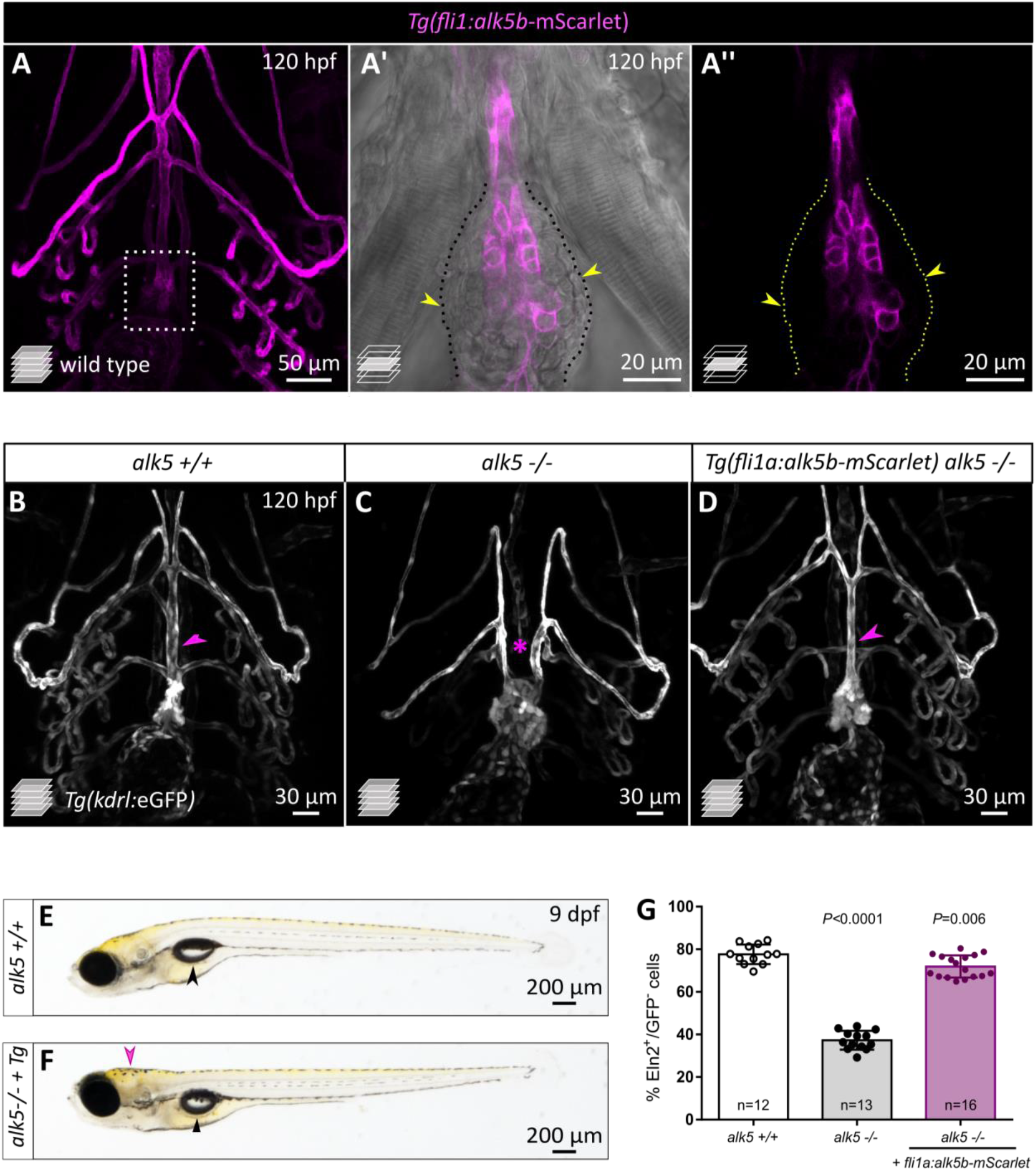
*alk5b* EC-specific overexpression restores the vascular network in *alk5* mutants – related to Figure 4. A-A’’) Ventral view of 120 hpf *Tg(fli1:alk5b-mScarlet)* larvae, showing the restricted expression of the transgene in the ECs. Boxed area shown in A’ and A’’; dotted lines outline the OFT; arrowheads, SMCs. B-D) Confocal images of the OFT in 120 hpf *Tg(kdrl:eGFP)* larvae, showing the formation of the VA in *alk5* mutants carrying the EC-specific rescue transgene (D, arrowheads) and its absence in *alk5* mutants (C, asterisk). E, F) Brightfield images of 9 dpf wild-type and *alk5* mutant animals carrying the EC-specific rescue transgene (*Tg*). Black arrowheads, swim bladder; pink arrowhead, deformation of the head. G) Quantification of the percentage of SMCs surrounded by Eln2 immunostaining (per sagittal plane) at 75 hpf. Means ± SD; *P* value from One way ANOVA test. Scale bars: A) 50 μm; A’, A’’) 20 μm; B, D) 30 μm; E, F) 200 μm.

**Figure S4 –.**
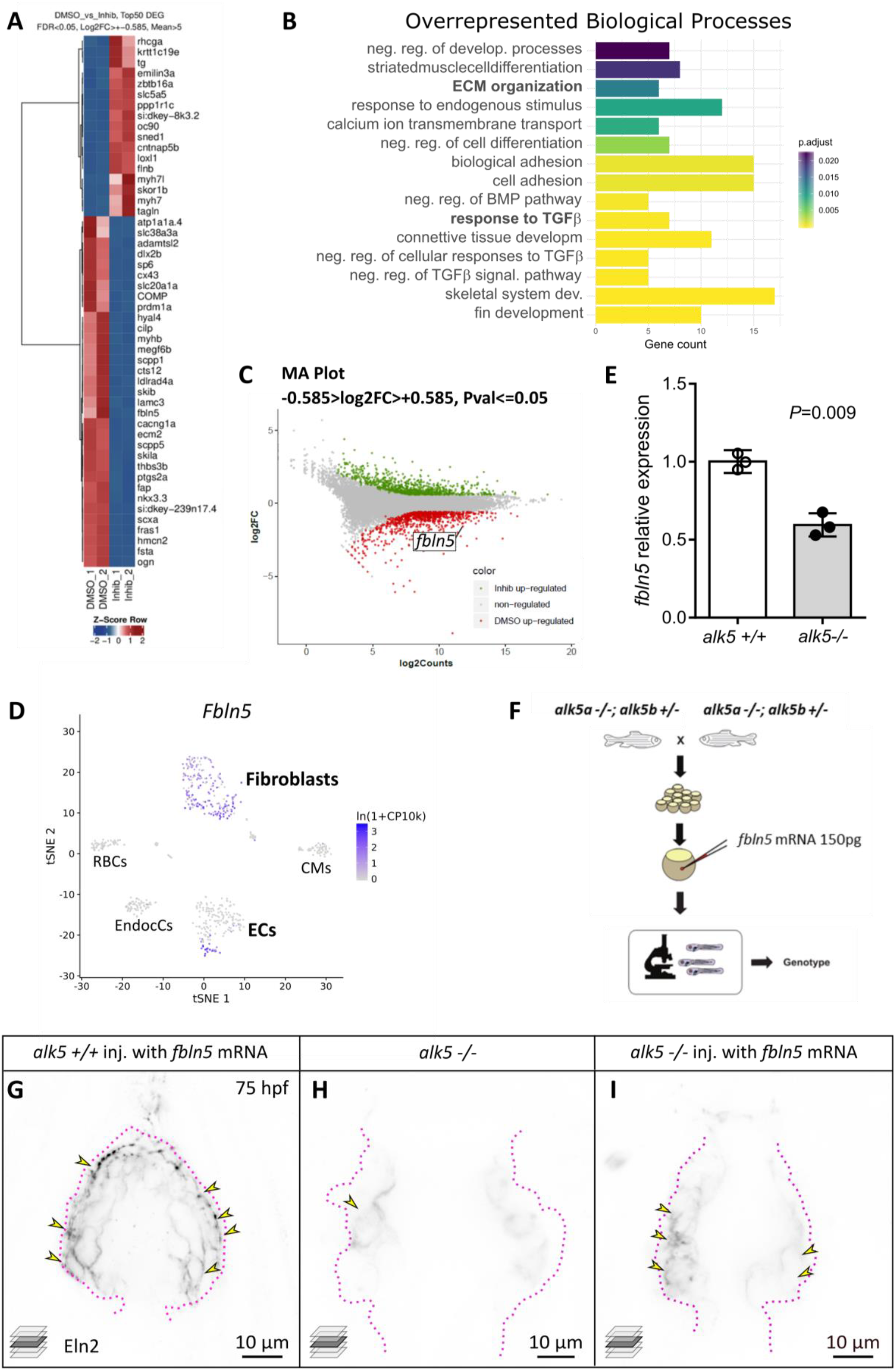
*Fbln5* is expressed in both SMCs and ECs in the adult mouse heart and it functions downstream of Alk5 in the zebrafish OFT-related to Figure 5. A) Heat map of the top 50 differentially expressed genes (DEGs) in Alk5 inhibitor treated embryonic hearts compared to controls. B) Gene ontology bar plot, showing some of the most overrepresented biological processes downregulated in the inhibitor treated embryos compared to control. C) MA plot displaying differentially expressed genes between inhibitor treated and control animals. *fbln5* is highlighted among the downregulated genes (red). D) Two-dimensional tSNE map showing the expression of *Fbln5* in different cells of heart and aorta in adult mice. Each cell is colored according to the scaled expression of *Fbln5* (Tabula Muris et al., 2018). E)*fbln5* relative mRNA levels in 72 hpf wild-type and *alk5* mutant larval hearts. Means ± SD; *P* values from *t*-tests. F) Schematic representing the strategy used for *fbln5* wild-type mRNA injections. G-I) Confocal images of 75 hpf larvae immunostained for Eln2 (black), following injection of *fbln5*. Yellow arrowheads, SMCs surrounded by Eln2 immunostaining; dotted lines outline the OFT. ECs, endothelial cells; EndocCs, endocardial cells; CMs, cardiomyocytes; RBCs, red blood cells. Scale bars: G-I) 10 μm.

## References

Albig, A.R., and Schiemann, W.P. (2004). Fibulin-5 antagonizes vascular endothelial growth factor (VEGF) signaling and angiogenic sprouting by endothelial cells. DNA Cell Biol 23, 367–79.

Anderson, R.H., Mori, S., Spicer, D.E., Brown, N.A., and Mohun, T.J. (2016). Development and Morphology of the Ventricular Outflow Tracts. World J Pediatr Congenit Heart Surg 7, 561–77.

Bochenek, M.L., Leidinger, C., Rosinus, N.S., Gogiraju, R., Guth, S., Hobohm, L., Jurk, K., Mayer, E., Munzel, T., Lankeit, M., et al. (2020). Activated Endothelial TGFbeta1 Signaling Promotes Venous Thrombus Nonresolution in Mice Via Endothelin-1: Potential Role for Chronic Thromboembolic Pulmonary Hypertension. Circ Res 126, 162–181.

Brownstein, A.J., Ziganshin, B.A., Kuivaniemi, H., Body, S.C., Bale, A.E., and Elefteriades, J.A. (2017). Genes Associated with Thoracic Aortic Aneurysm and Dissection: An Update and Clinical Implications. Aorta (Stamford) 5, 11–20.

Buckingham, M., Meilhac, S., and Zaffran, S. (2005). Building the mammalian heart from two sources of myocardial cells. Nat Rev Genet 6, 826–35.

Carvalho, R.L., Itoh, F., Goumans, M.J., Lebrin, F., Kato, M., Takahashi, S., Ema, M., Itoh, S., van Rooijen, M., Bertolino, P., et al. (2007). Compensatory signalling induced in the yolk sac vasculature by deletion of TGFbeta receptors in mice. J Cell Sci 120, 4269–77.

Chapman, S.L., Sicot, F.X., Davis, E.C., Huang, J., Sasaki, T., Chu, M.L., and Yanagisawa, H. (2010). Fibulin-2 and fibulin-5 cooperatively function to form the internal elastic lamina and protect from vascular injury. Arterioscler Thromb Vasc Biol 30, 68–74.

Choudhary, B., Zhou, J., Li, P., Thomas, S., Kaartinen, V., and Sucov, H.M. (2009). Absence of TGFbeta signaling in embryonic vascular smooth muscle leads to reduced lysyl oxidase expression, impaired elastogenesis, and aneurysm. Genesis 47, 115–21.

Cunha, S.I., Magnusson, P.U., Dejana, E., and Lampugnani, M.G. (2017). Deregulated TGF-beta/BMP Signaling in Vascular Malformations. Circ Res 121, 981–999.

Dave, J.M., Mirabella, T., Weatherbee, S.D., and Greif, D.M. (2018). Pericyte ALK5/TIMP3 Axis Contributes to Endothelial Morphogenesis in the Developing Brain. Dev Cell 44, 665–678 e6.

Davis, G.E., and Senger, D.R. (2005). Endothelial extracellular matrix: biosynthesis, remodeling, and functions during vascular morphogenesis and neovessel stabilization. Circ Res 97, 1093–107.

El-Brolosy, M.A., Kontarakis, Z., Rossi, A., Kuenne, C., Gunther, S., Fukuda, N., Kikhi, K., Boezio, G.L.M., Takacs, C.M., Lai, S.L., et al. (2019). Genetic compensation triggered by mutant mRNA degradation. Nature 568, 193–197.

Felker, A., Prummel, K.D., Merks, A.M., Mickoleit, M., Brombacher, E.C., Huisken, J., Panakova, D., and Mosimann, C. (2018). Continuous addition of progenitors forms the cardiac ventricle in zebrafish. Nat Commun 9, 2001.

Gaengel, K., Genove, G., Armulik, A., and Betsholtz, C. (2009). Endothelial-mural cell signaling in vascular development and angiogenesis. Arterioscler Thromb Vasc Biol 29, 630–8.

Gao, Y., Chen, T., and Raj, J.U. (2016). Endothelial and Smooth Muscle Cell Interactions in the Pathobiology of Pulmonary Hypertension. Am J Respir Cell Mol Biol 54, 451–60.

Gauvrit, S., Villasenor, A., Strilic, B., Kitchen, P., Collins, M.M., Marin-Juez, R., Guenther, S., Maischein, H.M., Fukuda, N., Canham, M.A., et al. (2018). HHEX is a transcriptional regulator of the VEGFC/FLT4/PROX1 signaling axis during vascular development. Nat Commun 9, 2704.

Gillis, E., Van Laer, L., and Loeys, B.L. (2013). Genetics of thoracic aortic aneurysm: at the crossroad of transforming growth factor-beta signaling and vascular smooth muscle cell contractility. Circ Res 113, 327–40.

Goumans, M.J., Lebrin, F., and Valdimarsdottir, G. (2003). Controlling the angiogenic switch: a balance between two distinct TGF-b receptor signaling pathways. Trends Cardiovasc Med 13, 301–7.

Goumans, M.J., and Ten Dijke, P. (2018). TGF-beta Signaling in Control of Cardiovascular Function. Cold Spring Harb Perspect Biol 10.

Goumans, M.J., Valdimarsdottir, G., Itoh, S., Rosendahl, A., Sideras, P., and ten Dijke, P. (2002). Balancing the activation state of the endothelium via two distinct TGF-beta type I receptors. EMBO J 21, 1743–53.

Grimes, A.C., and Kirby, M.L. (2009). The outflow tract of the heart in fishes: anatomy, genes and evolution. J Fish Biol 74, 983–1036.

Guner-Ataman, B., Paffett-Lugassy, N., Adams, M.S., Nevis, K.R., Jahangiri, L., Obregon, P., Kikuchi, K., Poss, K.D., Burns, C.E., and Burns, C.G. (2013). Zebrafish second heart field development relies on progenitor specification in anterior lateral plate mesoderm and nkx2.5 function. Development 140, 1353–63.

Guo, X., and Chen, S.Y. (2012). Transforming growth factor-beta and smooth muscle differentiation. World J Biol Chem 3, 41–52.

Hamm, M.J., Kirchmaier, B.C., and Herzog, W. (2016). Sema3d controls collective endothelial cell migration by distinct mechanisms via Nrp1 and PlxnD1. J Cell Biol 215, 415–430.

Jaffe, M., Sesti, C., Washington, I.M., Du, L., Dronadula, N., Chin, M.T., Stolz, D.B., Davis, E.C., and Dichek, D.A. (2012). Transforming growth factor-beta signaling in myogenic cells regulates vascular morphogenesis, differentiation, and matrix synthesis. Arterioscler Thromb Vasc Biol 32, e1–11.

Jana, S., Hu, M., Shen, M., and Kassiri, Z. (2019). Extracellular matrix, regional heterogeneity of the aorta, and aortic aneurysm. Exp Mol Med 51, 160.

Kelleher, C.M., McLean, S.E., and Mecham, R.P. (2004). Vascular extracellular matrix and aortic development. Curr Top Dev Biol 62, 153–88.

Kelly, R.G., and Buckingham, M.E. (2002). The anterior heart-forming field: voyage to the arterial pole of the heart. Trends Genet 18, 210–6.

Kim, H.W., and Stansfield, B.K. (2017). Genetic and Epigenetic Regulation of Aortic Aneurysms. Biomed Res Int 2017, 7268521.

Knight, H.G., and Yelon, D. (2016). Utilizing Zebrafish to Understand Second Heart Field Development. In Etiology and Morphogenesis of Congenital Heart Disease: From Gene Function and Cellular Interaction to Morphology, Nakanishi, T., Markwald, R.R., Baldwin, H.S., Keller, B.B., Srivastava, D. and Yamagishi, H., ed. (Tokyo, pp. 193–199.

Larsson, J., Goumans, M.J., Sjostrand, L.J., van Rooijen, M.A., Ward, D., Leveen, P., Xu, X., ten Dijke, P., Mummery, C.L., and Karlsson, S. (2001). Abnormal angiogenesis but intact hematopoietic potential in TGF-beta type I receptor-deficient mice. EMBO J 20, 1663–73.

Lebrin, F., Goumans, M.J., Jonker, L., Carvalho, R.L., Valdimarsdottir, G., Thorikay, M., Mummery, C., Arthur, H.M., and ten Dijke, P. (2004). Endoglin promotes endothelial cell proliferation and TGF-beta/ALK1 signal transduction. EMBO J 23, 4018–28.

Li, M., Qian, M., Kyler, K., and Xu, J. (2018). Endothelial-Vascular Smooth Muscle Cells Interactions in Atherosclerosis. Front Cardiovasc Med 5, 151.

Li, W., Li, Q., Jiao, Y., Qin, L., Ali, R., Zhou, J., Ferruzzi, J., Kim, R.W., Geirsson, A., Dietz, H.C., et al. (2014). Tgfbr2 disruption in postnatal smooth muscle impairs aortic wall homeostasis. J Clin Invest 124, 755–67.

Lilly, B. (2014). We have contact: endothelial cell-smooth muscle cell interactions. Physiology (Bethesda) 29, 234–41.

Liu, X., Chen, W., Li, W., Li, Y., Priest, J.R., Zhou, B., Wang, J., and Zhou, Z. (2019). Single-Cell RNA-Seq of the Developing Cardiac Outflow Tract Reveals Convergent Development of the Vascular Smooth Muscle Cells. Cell Rep 28, 1346–1361 e4.

Mancini, M.L., Terzic, A., Conley, B.A., Oxburgh, L.H., Nicola, T., and Vary, C.P. (2009). Endoglin plays distinct roles in vascular smooth muscle cell recruitment and regulation of arteriovenous identity during angiogenesis. Dev Dyn 238, 2479–93.

Maring, J.A., van Meeteren, L.A., Goumans, M.J., and Ten Dijke, P. (2016). Interrogating TGF-beta Function and Regulation in Endothelial Cells. Methods Mol Biol 1344, 193203.

Massague, J. (2012). TGFbeta signalling in context. Nat Rev Mol Cell Biol 13, 616–30.

Miao, M., Bruce, A.E., Bhanji, T., Davis, E.C., and Keeley, F.W. (2007). Differential expression of two tropoelastin genes in zebrafish. Matrix Biology 26, 115–24.

Mullapudi, S.T., Helker, C.S., Boezio, G.L., Maischein, H.M., Sokol, A.M., Guenther, S., Matsuda, H., Kubicek, S., Graumann, J., Yang, Y.H.C., et al. (2018). Screening for insulin-independent pathways that modulate glucose homeostasis identifies androgen receptor antagonists. Elife 7.

Nakamura, T., Lozano, P.R., Ikeda, Y., Iwanaga, Y., Hinek, A., Minamisawa, S., Cheng, C.F., Kobuke, K., Dalton, N., Takada, Y., et al. (2002). Fibulin-5/DANCE is essential for elastogenesis in vivo. Nature 415, 171–5.

Nakamura, T., Ruiz-Lozano, P., Lindner, V., Yabe, D., Taniwaki, M., Furukawa, Y., Kobuke, K., Tashiro, K., Lu, Z., Andon, N.L., et al. (1999). DANCE, a novel secreted RGD protein expressed in developing, atherosclerotic, and balloon-injured arteries. J Biol Chem 274, 22476–83.

Nauroy, P., Hughes, S., Naba, A., and Ruggiero, F. (2018). The in-silico zebrafish matrisome: A new tool to study extracellular matrix gene and protein functions. Matrix Biol 65, 5–13.

Neeb, Z., Lajiness, J.D., Bolanis, E., and Conway, S.J. (2013). Cardiac outflow tract anomalies. Wiley Interdiscip Rev Dev Biol 2, 499–530.

Orriols, M., Varona, S., Marti-Pamies, I., Galan, M., Guadall, A., Escudero, J.R., Martin-Ventura, J.L., Camacho, M., Vila, L., Martinez-Gonzalez, J., et al. (2016). Down-regulation of Fibulin-5 is associated with aortic dilation: role of inflammation and epigenetics. Cardiovasc Res 110, 431–42.

Paffett-Lugassy, N., Novikov, N., Jeffrey, S., Abrial, M., Guner-Ataman, B., Sakthivel, S., Burns, C.E., and Burns, C.G. (2017). Unique developmental trajectories and genetic regulation of ventricular and outflow tract progenitors in the zebrafish second heart field. Development 144, 4616–4624.

Pardali, E., Goumans, M.J., and ten Dijke, P. (2010). Signaling by members of the TGF-beta family in vascular morphogenesis and disease. Trends Cell Biol 20, 556–67.

Pauli, A., Valen, E., Lin, M.F., Garber, M., Vastenhouw, N.L., Levin, J.Z., Fan, L., Sandelin, A., Rinn, J.L., Regev, A., et al. (2012). Systematic identification of long noncoding RNAs expressed during zebrafish embryogenesis. Genome Res 22, 577–91.

Perbellini, F., Watson, S.A., Bardi, I., and Terracciano, C.M. (2018). Heterocellularity and Cellular Cross-Talk in the Cardiovascular System. Front Cardiovasc Med 5, 143.

Perrucci, G.L., Rurali, E., Gowran, A., Pini, A., Antona, C., Chiesa, R., Pompilio, G., and Nigro, P. (2017). Vascular smooth muscle cells in Marfan syndrome aneurysm: the broken bricks in the aortic wall. Cell Mol Life Sci 74, 267–277.

Preis, M., Cohen, T., Sarnatzki, Y., Ben Yosef, Y., Schneiderman, J., Gluzman, Z., Koren, B., Lewis, B.S., Shaul, Y., and Flugelman, M.Y. (2006). Effects of fibulin-5 on attachment, adhesion, and proliferation of primary human endothelial cells. Biochem Biophys Res Commun 348, 1024–33.

Raines, E.W. (2000). The extracellular matrix can regulate vascular cell migration, proliferation, and survival: relationships to vascular disease. Int J Exp Pathol 81, 173–82.

Rochon, E.R., Menon, P.G., and Roman, B.L. (2016). Alk1 controls arterial endothelial cell migration in lumenized vessels. Development 143, 2593–602.

Segers, V.F.M., Brutsaert, D.L., and De Keulenaer, G.W. (2018). Cardiac Remodeling: Endothelial Cells Have More to Say Than Just NO. Front Physiol 9, 382.

Seki, T., Hong, K.H., and Oh, S.P. (2006). Nonoverlapping expression patterns of ALK1 and ALK5 reveal distinct roles of each receptor in vascular development. Lab Invest 86, 116–29.

Singh, A.R., Sivadas, A., Sabharwal, A., Vellarikal, S.K., Jayarajan, R., Verma, A., Kapoor, S., Joshi, A., Scaria, V., and Sivasubbu, S. (2016). Chamber Specific Gene Expression Landscape of the Zebrafish Heart. PLoS One 11, e0147823.

Sridurongrit, S., Larsson, J., Schwartz, R., Ruiz-Lozano, P., and Kaartinen, V. (2008). Signaling via the Tgf-beta type I receptor Alk5 in heart development. Dev Biol 322, 208–18.

Stainier, D.Y., and Fishman, M.C. (1994). The zebrafish as a model system to study cardiovascular development. Trends Cardiovasc Med 4, 207–12.

Stratman, A.N., Pezoa, S.A., Farrelly, O.M., Castranova, D., Dye, L.E., 3rd, Butler, M.G., Sidik, H., Talbot, W.S., and Weinstein, B.M. (2017). Interactions between mural cells and endothelial cells stabilize the developing zebrafish dorsal aorta. Development 144, 115–127.

Sugishita, Y., Watanabe, M., and Fisher, S.A. (2004). The development of the embryonic outflow tract provides novel insights into cardiac differentiation and remodeling. Trends Cardiovas Med 14, 235–241.

Sullivan, K.M., Bissonnette, R., Yanagisawa, H., Hussain, S.N., and Davis, E.C. (2007). Fibulin-5 functions as an endogenous angiogenesis inhibitor. Lab Invest 87, 818–27.

Sun, J., Deng, H., Zhou, Z., Xiong, X., and Gao, L. (2018). Endothelium as a Potential Target for Treatment of Abdominal Aortic Aneurysm. Oxid Med Cell Longev 2018, 6306542.

Sweeney, M., and Foldes, G. (2018). It Takes Two: Endothelial-Perivascular Cell Cross-Talk in Vascular Development and Disease. Front Cardiovasc Med 5, 154.

Tabula Muris, C., Overall, c., Logistical, c., Organ, c., processing, Library, p., sequencing, Computational data, a., Cell type, a., Writing, g., et al. (2018). Single-cell transcriptomics of 20 mouse organs creates a Tabula Muris. Nature 562, 367–372.

Takeda, N., Hara, H., Fujiwara, T., Kanaya, T., Maemura, S., and Komuro, I. (2018). TGF-beta Signaling-Related Genes and Thoracic Aortic Aneurysms and Dissections. Int J Mol Sci 19.

Todorovic, V., Frendewey, D., Gutstein, D.E., Chen, Y., Freyer, L., Finnegan, E., Liu, F., Murphy, A., Valenzuela, D., Yancopoulos, G., et al. (2007). Long form of latent TGF-beta binding protein 1 (Ltbp1L) is essential for cardiac outflow tract septation and remodeling. Development 134, 3723–32.

van de Pol, V., Kurakula, K., DeRuiter, M.C., and Goumans, M.J. (2017). Thoracic Aortic Aneurysm Development in Patients with Bicuspid Aortic Valve: What Is the Role of Endothelial Cells? Front Physiol 8, 938.

van Meeteren, L.A., and ten Dijke, P. (2012). Regulation of endothelial cell plasticity by TGF-beta. Cell Tissue Res 347, 177–86.

Waldo, K.L., Hutson, M.R., Stadt, H.A., Zdanowicz, M., Zdanowicz, J., and Kirby, M.L. (2005a). Cardiac neural crest is necessary for normal addition of the myocardiurn to the arterial pole from the secondary heart field. Dev Biol 281, 66–77.

Waldo, K.L., Hutson, M.R., Ward, C.C., Zdanowicz, M., Stadt, H.A., Kumiski, D., Abu-Issa, R., and Kirby, M.L. (2005b). Secondary heart field contributes myocardium and smooth muscle to the arterial pole of the developing heart. Dev Biol 281, 78–90.

Williamson, M.R., Shuttleworth, A., Canfield, A.E., Black, R.A., and Kielty, C.M. (2007). The role of endothelial cell attachment to elastic fibre molecules in the enhancement of monolayer formation and retention, and the inhibition of smooth muscle cell recruitment. Biomaterials 28, 5307–18.

Yanagisawa, H., Schluterman, M.K., and Brekken, R.A. (2009). Fibulin-5, an integrin-binding matricellular protein: its function in development and disease. J Cell Commun Signal 3, 337–47.

Yang, H., Zhou, Y., Gu, J., Xie, S., Xu, Y., Zhu, G., Wang, L., Huang, J., Ma, H., and Yao, J. (2013). Deep mRNA sequencing analysis to capture the transcriptome landscape of zebrafish embryos and larvae. PLoS One 8, e64058.

Yang, P., Schmit, B.M., Fu, C., DeSart, K., Oh, S.P., Berceli, S.A., and Jiang, Z. (2016). Smooth muscle cell-specific Tgfbr1 deficiency promotes aortic aneurysm formation by stimulating multiple signaling events. Sci Rep 6, 35444.

Zhang, P., Hou, S., Chen, J., Zhang, J., Lin, F., Ju, R., Cheng, X., Ma, X., Song, Y., Zhang, Y., et al. (2016). Smad4 Deficiency in Smooth Muscle Cells Initiates the Formation of Aortic Aneurysm. Circ Res 118, 388–99.

Zhang, Y.E. (2018). Mechanistic insight into contextual TGF-beta signaling. Curr Opin Cell Biol 51, 1–7.

Zhou, Y., Cashman, T.J., Nevis, K.R., Obregon, P., Carney, S.A., Liu, Y., Gu, A., Mosimann, C., Sondalle, S., Peterson, R.E., et al. (2011). Latent TGF-beta binding protein 3 identifies a second heart field in zebrafish. Nature 474, 645–8.

